# FGF1 Promotes *Xenopus laevis* Lens Regeneration

**DOI:** 10.1101/411991

**Authors:** Lisa Moore, Kimberly J. Perry, Cindy Sun, Jonathan J. Henry

## Abstract

**Background:** The frog *Xenopus laevis* has notable regenerative capabilities, including that of the lens. The neural retina provides the factors that trigger lens regeneration from the cornea, but the identity of these factors is largely unknown. In contrast to the cornea, fibroblast growth factors FGF1, 8, and 9 are highly expressed within the retina, and are potential candidates for those factors. The purpose of this study is to determine whether specific FGF proteins can induce lens formation, and if perturbation of FGFR signaling inhibits lens regeneration.

**Methods:** A novel cornea epithelial culture method was developed to investigate the sufficiency of FGFs in lens regeneration. Additionally, transgenic larvae expressing dominant negative FGFR1 were used to investigate the necessity of FGFR signaling in lens regeneration.

**Results:** Treatment of cultured corneas with FGF1 induced lens regeneration in a dose-dependent manner, whereas treatment with FGF2, FGF8, or FGF9 did not result in significant lens regeneration. Inhibition of FGFR signaling decreased the lens regeneration rate for *in vitro* eye cultures.

**Conclusion:** The culture techniques developed here, and elsewhere, have provided reliable methods for examining the necessity of various factors that may be involved in lens regeneration. Based on the results demonstrated in this study, we found that FGF1 signaling and FGFR activation are key factors for lens regeneration in *Xenopus*.

## Introduction

Lens regeneration is a unique phenomenon among some vertebrates. Wolffian lens regeneration is one process of lens regeneration, which occurs among newts and salamanders (Colluci, 1981; Wolff, 1895). In those animals the dorsal pigmented iris undergoes transdifferentiation to form a lens (Henry, 2003; Henry & Tsonis, 2010; Barbosa-Sabanero et al., 2012). The frog *Xenopus* can also regenerate a new lens following removal of the original lens, but this regenerated tissue is derived instead from the basal layer of the cornea epithelium (Freeman, 1963). Experimental evidence indicates that *Xenopus* lens regeneration is initiated when diffusible factors from the neural retina are permitted to reach the cornea (Bosco et al., 1979, 1980, 1997a).

The fibroblast growth factor (FGF) pathway is one group of factors that appear to play a key role for lens regeneration in these different systems. Culture experiments including exogenous FGFs, suggest that FGFs may be sufficient to trigger lens regeneration (Bosco et al., 1997b; Hayashi et al., 2002, 2004). For example, in newts, FGF2 treatment has been shown to be sufficient for inducing Wolffian lens formation both *in vitro* using dorsal iris cell reaggregates (Hayashi et al., 2002), and *in vivo* upon injection into the anterior or posterior eye chamber (Hayashi et al., 2004). Treatment with high levels of FGF1 appears to induce lens cell formation in explanted *Xenopus* cornea tissues (Bosco et al., 1997b; Bosco 1998). Similarly, injection of FGF1 into lentectomized adult newt eyes results in Wolffian lens formation from only the dorsal iris and not the ventral iris, but the formed lenses have abnormal fiber cells, and a thinned or absent lens epithelium (Yang et al., 2005). Additionally, the necessity for FGFR activity during lens regeneration has been demonstrated by SU5402 small molecule inhibitor experiments in both Wolffian lens regeneration (Del Rio-Tsonis et al., 1998), and *Xenopus* lens regeneration, although SU5402 can also inhibit other receptor tyrosine kinases. Expression analyses suggest that FGF1, FGF8, and FGF9 could play roles in *Xenopus* lens regeneration (Fukui and Henry, 2001), as these factors are robustly expressed in the retina and not the cornea. Neither FGF2, FGF8 nor FGF9 have been tested previously for their potential to induce lens formation in *Xenopus* cornea tissue.

To undertake these experiments, we developed a novel culture method that enables corneal tissue to maintain a flattened epithelial morphology. Using this new method, this paper investigates whether FGF1, FGF2, FGF8, or FGF9 are sufficient to induce lens regeneration in cultured *Xenopus* corneas. We found that FGF1, and to a much lesser extent, FGF8, triggered lens cell differentiation. Additionally, we examined the necessity of FGFR signaling for lens regeneration by conditional expression of a dominant negative FGFR1 in transgenic eye tissues. Heat shock activated expression of dominant negative FGFR1 (XFD) indicated that FGFR function is important for lens regeneration. These results provide evidence that the FGF pathway is both necessary and sufficient for the regenerative process.

## Materials and Methods

### Cornea culture

Cornea epithelial tissues from larval tadpoles (stages 51-54; Nieuwkoop and Faber, 1956) were cultured alone without the retina (**Figure 1**). Corneas were excised from wild-type larvae using ultrafine scissors and transferred to a wash dish containing culture medium and finally to a 24-well plate well. The wells were coated with gelatin (30 minute incubation with autoclaved 0.1% Knox unflavored gelatin, Kraft foods, Northfield, IL) to promote cellular attachment to the bottom of the well. The corneal tissue was oriented so the inner basal layer was in contact with the gelatin-coated bottom surface of the culture well, and was affixed to the bottom of the well by applying firm pressure with the sharpened tips of #5 Dumont forceps. This pressure created slight indentations in the plastic culture well, effectively “pinning” the corneas in place (**Figure 1B-C**). The neural retina, the natural source of lens regeneration factors, was omitted in these cultures to allow for assaying whether specific FGFs may be sufficient to induce lens regeneration. The serum free, 67% L-15 culture medium (Invitrogen, Grand Island, NY) contained Pen-Strep (100 U/ml Penicillin and 100 μg/ml Streptomycin, Mediatech, Herndon, VA), marbofloxacin (10 μg/ml, Sigma-Aldrich), and Amphotericin B (2.5 μg/ml, Sigma-Aldrich) to inhibit bacterial and/or fungal growth. Four corneas were individually attached in each well and the media was changed every three days. Corneas cultured using this method adhere to the bottom of the well and start spreading within two days (**Figure 1C-D**). Corneal cultures were treated with recombinant human FGF1, FGF2, FGF8, and FGF9 (R&D Systems, Minneapolis, MN, catalog numbers 232-FA-025, 233-FB-025, 423-F8-025, and 273-F9-025, respectively) at various concentrations indicated in **Table 1**. Control cultures were treated with an equivalent dilution of the stock FGF carrier (PBS with 0.1% BSA) as used for the highest concentrations of the FGFs tested. After 9 days, the cornea cultures were fixed in 3.7% formaldehyde, and antibody staining was performed to detect *Xenopus* lens proteins with a rabbit anti-lens polyclonal antibody (**Figures 2-3**; Henry and Grainger, 1990). The specificity of the anti-lens polyclonal antibody has been verified previously (Henry and Grainger, 1990) and used in other studies (Fukui & Henry, 2001; Walter et al., 2008; Perry et al., 2010, 2013; Thomas & Henry, 2014). Corneas remained in the wells for fixation, staining and imaging steps. Statistical analysis was performed as described below.

**Table 1:** The formation of lens structures was quantified for *in vitro* cornea cultures treated with FGF1, FGF2, FGF8 or FGF9. The respective concentrations of FGFs tested, the number of cases forming anti-lens antibody positive, lens cell aggregates, and total numbers of corneas assayed are shown for each condition, and p-values are included for each corresponding data point. Positive lens cell formation was determined using anti-lens antibody staining after FGF treatment for 9 days in culture. Statistical analysis was performed using Fisher’s exact test to compare controls with each experimental condition.

|  | Concentration | + Regeneration | Total Corneas | p-Value |
| --- | --- | --- | --- | --- |
| FGF1 | control | 1 (3%) | 36 |  |
|  | 10ng/ml | 1 (4%) | 24 | 0.644 |
|  | 20ng/ml | 4 (19%) | 21 | 0.0563 |
|  | 50ng/ml | 6 (26%) | 23 | 0.0114 |
|  | 100ng/ml | 7 (41%) | 17 | 0.0075 |
|  | 500ng/ml | 9 (47%) | 19 | 0.00012 |
|  | 1ug/ml | 10 (53%) | 19 | 0.000028 |
| FGF2 | control | 1 (6%) | 16 |  |
|  | 10ng/ml | 0 (0%) | 16 | 1 |
|  | 20ng/ml | 0 (0%) | 16 | 1 |
|  | 50ng/ml | 1 (7%) | 14 | 0.7241 |
|  | 100ng/ml | 0 (0%) | 16 | 1 |
|  | 500ng/ml | 0 (0%) | 16 | 1 |
| FGF8 | control | 0 (0%) | 15 |  |
|  | 10ng/ml | 0 (0%) | 15 | 1 |
|  | 20ng/ml | 0 (0%) | 15 | 1 |
|  | 50ng/ml | 0 (0%) | 15 | 1 |
|  | 100ng/ml | 1 (7%) | 15 | 0.5 |
|  | 500ng/ml | 3 (19%) | 16 | 0.1246 |
| FGF9 | control | 1 (6%) | 16 |  |
|  | 10ng/ml | 0 (0%) | 15 | 1 |
|  | 20ng/ml | 0 (0%) | 13 | 1 |
|  | 50ng/ml | 1 (8%) | 13 | 1 |
|  | 100ng/ml | 1 (7%) | 14 | 0.7241 |
|  | 500ng/ml | 0 (0%) | 16 | 1 |

**Figure 1:**
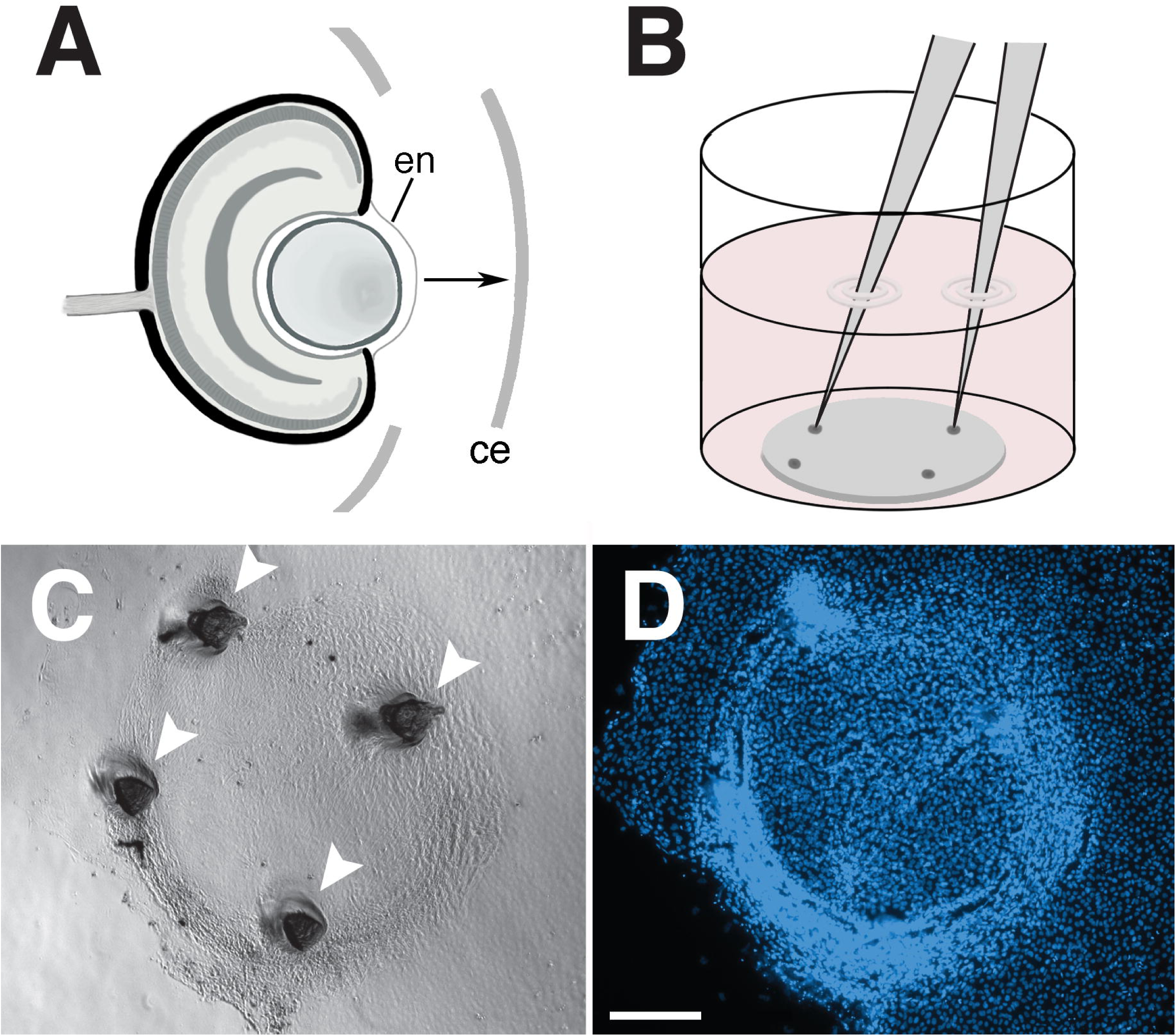
Cornea culture method to assay factors sufficient for lens formation. (A) The *Xenopus* cornea epithelium from a larval tadpole (st 51-54) is excised. During this procedure, the cornea endothelium of the donor eye remains intact and attached to the eye cup, preventing any lens tissue from contaminating the explanted cornea epithelial culture. (B) Diagram of the method used to attach the cornea to the bottom of the culture well. Gentle pressure is applied with sharpened forceps. (C, D) Cultured *Xenopus* cornea epithelium imaged using bright field illumination (C) and corresponding fluorescence detection of Hoescht nuclear stain (D). ce, cornea epithelium; en, cornea endothelium. White arrowheads in C indicate depressions in the plastic culture well created by the forceps. The scale bar in D indicates 250 μm.

**Figure 2:**
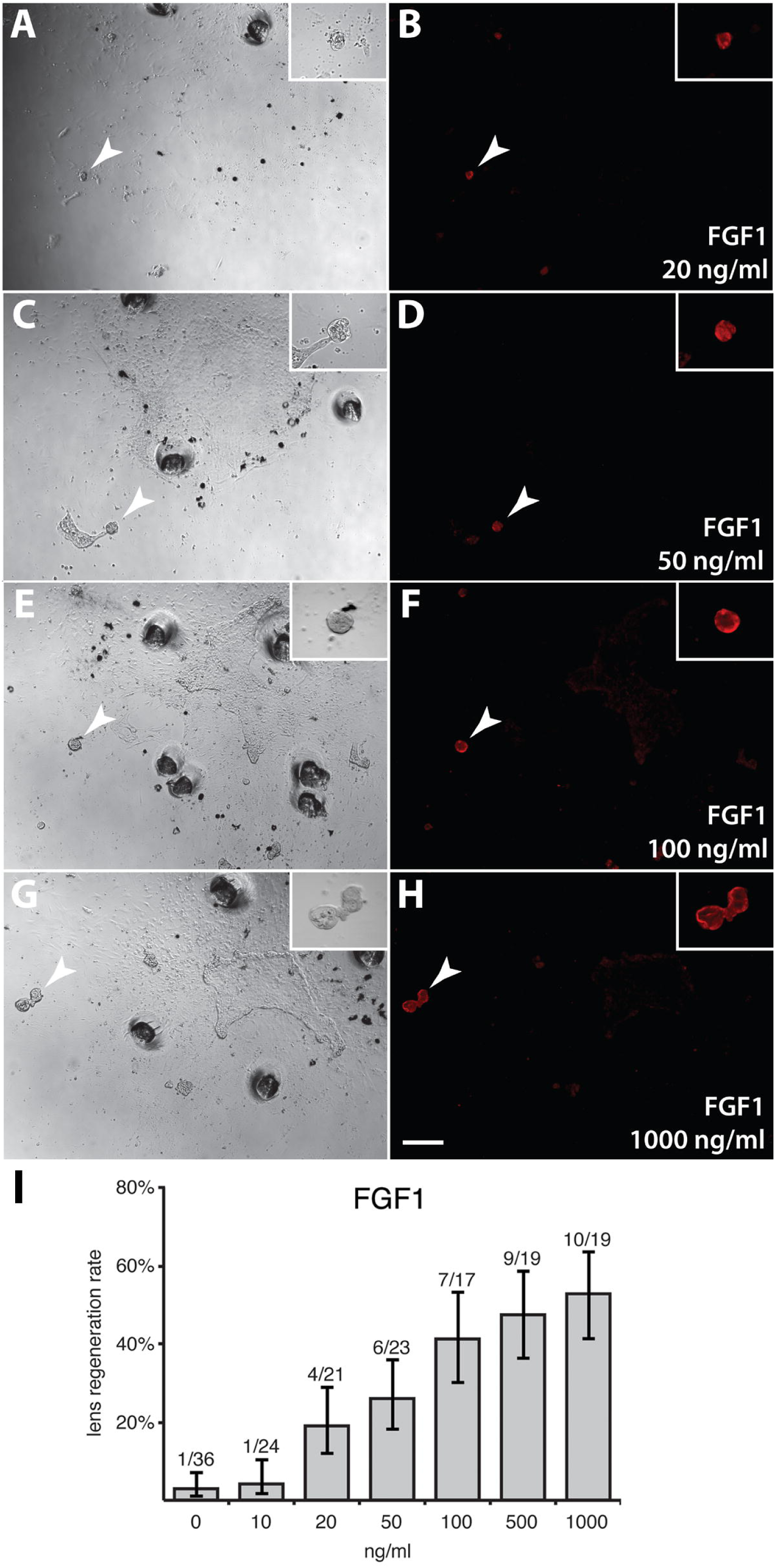
Corneas cultured for 9 days with various concentrations of FGF1 showing anti-lens antibody labeling. Corneas were treated with 20 ng/ml (A-B), 50 ng/ml (C-D), 100 ng/ml (E-F), 1000 ng/ml FGF1 (G-H). Images were taken under bright field (A, C, E, G) and fluorescence illumination to detect anti-lens antibody staining (B, D, F, H). Insets in A-H contain higher magnitude images of corresponding lenses in each panel. Depressions in the culture wells (dark circular spots in A, C, E, G) indicate points (indentations) where the cornea was secured to the bottom of the well using sharpened forceps. White arrowheads in A-H point to regenerated lenses. The scale bar in H indicates 100 μm for the insets in A-H, and 200 μm for panels A-H. The quantification of lens structure formation rates for FGF1 is shown in I. The leftmost bar (0 ng/ml) indicates the control condition in which no FGF was applied. The number of corneas forming lens cell aggregates and total number of corneas observed for each condition are indicated above each bar. The Y-axis indicates the % regeneration rate for each condition. Error bars show Wilson score confidence intervals where Z=1.

### FGF activity assay *in vitro*

Prior to testing on cornea cultures, the FGF proteins (1, 2, 8, 9) were each assayed in *Xenopus* eye cultures to determine whether they were capable of enhancing cell proliferation. Whole eyes were excised from *Xenopus laevis* wild-type larvae (stages 51-54), washed in serum free 67% L-15 media with Pen-Strep, marbofloxacin and Amphotericin B (concentrations mentioned above), and cultured in a 24-well plate. Each well contained four eyes in 0.5mL of culture media and treated eyes were cultured with FGF1, FGF2, FG8 or FGF9 (100ng/ml for each). Culture media containing each specific FGF protein was changed daily and eyes were cultured for 72 hours and then fixed in 3.7% formaldehyde. Antibody staining was performed on the whole eye with anti-Histone H3S10P antibody (1:400, sc-8656-R, Santa Cruz Biotechnology, Inc., Dallas, TX) to detect transcriptionally active chromatin. Following antibody labeling, corneas were removed from the eye cup and the cornea epithelium was placed on a slide for fluorescent imaging and analysis. Nuclei were labeled with DAPI and a standardized area was analyzed for each cornea. In this area total nuclei as well as histone labeled nuclei were counted, and the proportion of histone labeled nuclei to total nuclei was recorded for each sample for comparison to control, untreated eyes (**Figure 4**).

**Figure 4:**
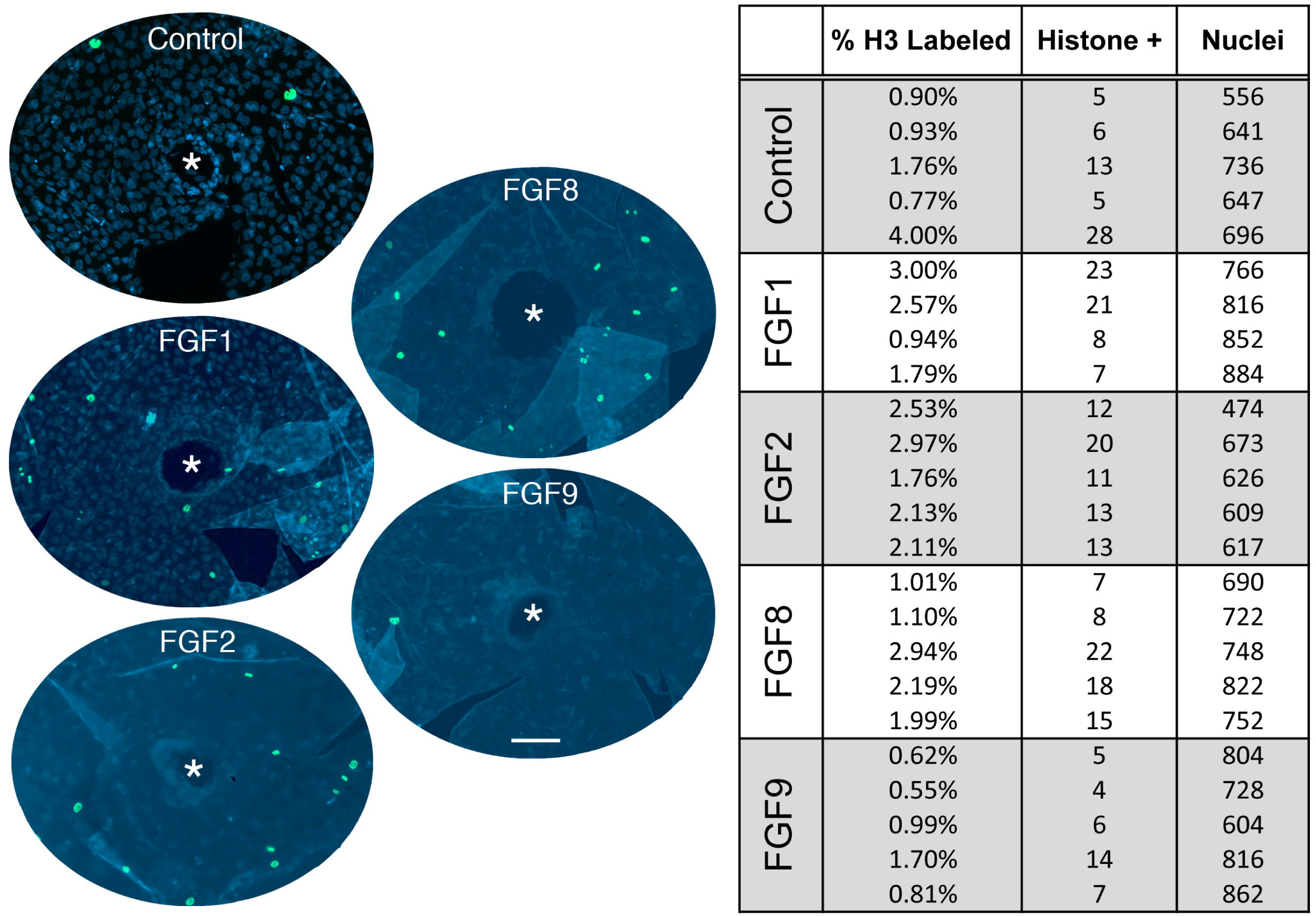
Results of protein activity experiments on representative corneas treated with FGF1, 2, 8 or 9. To confirm increased cell activity, the corneas were labeled with anti-Histone H3S10P antibody (green). Cell nuclei are shown in blue (DAPI-labeled). Increased chromatin labeling was observed for FGF1 (25% increase in histone labeled cells), FGF 2 (39% increase) and FGF8 (11% increase). Asterisk notes the center of the cornea and the oval area shown represents the area that was used for counting labeled nuclei. The scale bar in FGF9 indicates 50 μm.

### Plasmid construction

The transgenesis plasmid used here is based on the “heat shock green-eyed monster” (HGEM) construct (Beck et al., 2003). Here the plasmid (CGHG) contains an hsp70 promoter expressing the inserted transgene with a C-terminal GFP fusion, and a lens-specific gamma-crystallin promoter expressing GFP (**Figure 5**). Transgenic animals can be readily identified by the presence of green fluorescent lenses that express GFP (**Figure 5D-F**). We modified the CGHG vector by removing GFP (fused to hsp70) and inserted an HA recognition sequence (YPYDVPDYA) in its place (**Figure 5B**). An in-frame initiation codon upstream of the HA recognition sequence was included to ensure translation of the HA protein. Control transgenic animals were produced using this HA transgene, where HA was expressed following heat shock and was detectable with an anti-HA antibody (sc-805, Santa Cruz). In order to test the necessity of FGFR function in lens regeneration, a dominant negative FGFR1 (XFD) lacking the intracellular kinase domains was expressed to inhibit FGF downstream pathway activation (Amaya et al., 1991). *XFD* was inserted downstream of the hsp70 promoter and in-frame with the C-terminal HA tag (**Figure 5C**). This allowed heat-shock inducible expression of XFD (**Figure 5G**).

**Figure 5:**
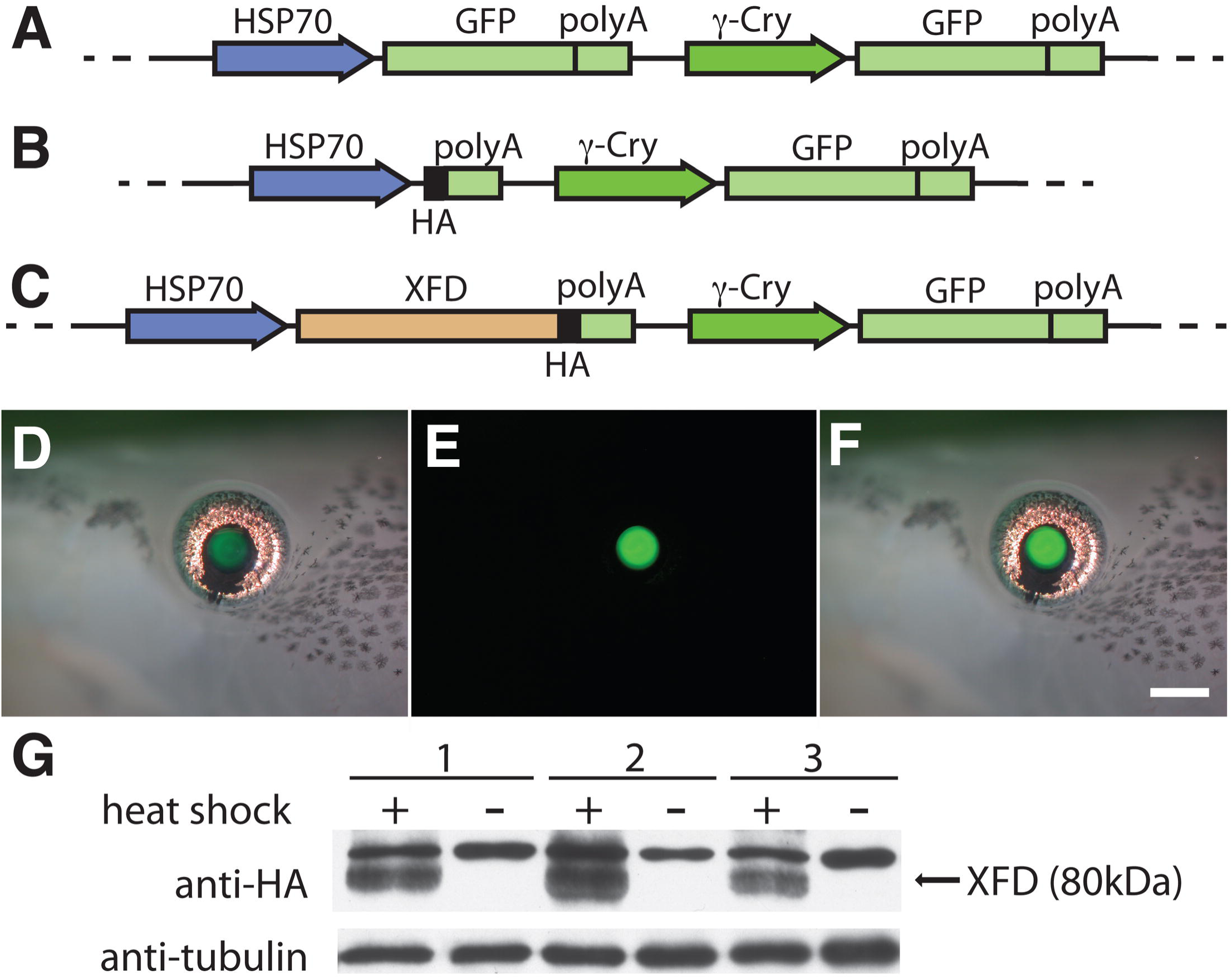
Diagrams of transgenesis plasmids used to explore the effect of FGFR inhibition on lens regeneration, and resulting transgenic larvae expressing GFP in lens tissue and XFD protein upon heat shock activation. (A) The original CGHG transgenesis construct containing the GFP reporters under HSP70 and gamma crystallin (γ-Cry) promoters. (B) The control plasmid, constructed from the original CGHG plasmid, now containing a heat shock promoter expressing only the HA tag. (C) The experimental transgenesis construct containing the dominant negative FGFR1 gene (XFD) expressed using the HSP70 heat shock promoter together with gamma-crystallin promoter driving expression of GFP (see text for further details). (D) An example of a transgenic larval eye under bright field illumination, noting that green fluorescence of the lens can be observed under visible light. (E) Corresponding fluorescence image of transgenic larval eye. (F) Overlay of fluorescence and visible images shown in D and E. (G) Expression of the XFD transgene was confirmed by Western blot analysis shown here for three specimens. Expression of the XFD protein containing a C-terminal HA tag was verified by anti-HA antibody (80 kDa). The antibody also detects an unknown higher molecular weight band (95 kDa) in all samples and anti-β-tubulin antibody was used as a protein loading control (55 kDa). GFP, sequence encoding green fluorescent protein; HA, hemagglutinin tag; polyA, poly(A) tail. The scale bar in F indicates 500 μm.

### Generation of transgenic *Xenopus laevis*

Adult *Xenopus laevis* animals were obtained from eNasco (Fort Atkinson, WI). Sperm nuclei for transgenesis were prepared as previously described (Sive et al., 2000). Transgenic larvae were generated using a protocol similar to the REMI method with slight modifications noted (Kroll et al., 1996; Hamilton et al., 2016). Sperm nuclei were incubated with linearized DNA without egg extract, as described in (Sparrow et al., 2000). Embryos were incubated in 0.1x Marc’s Modified Ringer (MMR) solution without ficoll starting at the four-cell stage. Transgenic larvae were identified by their green fluorescent lenses, indicative of *GFP* transgene expression, 4-6 days after fertilization (**Figure 5D-F**). Larvae were reared to various stages, as described previously (Henry & Grainger, 1987; Schaefer et al., 1999). In the case of older larvae (stage 47-54) used in the experiments below, the fluorescence was typically bright enough to observe green color in the eye using only ambient light (**Figure 5D-F**), but all cases were verified with fluorescence illumination.

### Transgenic *in vitro* eye culture

Corneas of transgenic larvae were cultured with retinas from isolated wild type larvae (**Figures 5-6**). As shown in the past, combining a neural retina with a cornea in culture is sufficient to recapitulate lens regeneration in *Xenopus* (Bosco et al., 1993c; Fukui & Henry, 2001). Cornea epithelia were excised from transgenic larvae (stages 49-55) using ultrafine scissors, and transferred to a wash dish with culture medium (modified L-15 medium containing fetal bovine serum (Kay and Peng, 1991), plus antibiotics/antifungals mentioned above). To prepare eye cups, lenses and corneas were first removed from wild type larval eyes (stages 50-55), and then the remaining retinal tissues were excised and transferred to the same wash dish containing the isolated corneas. A single transgenic cornea was gently inserted into the vitreous chamber of a corresponding wild type eye cup. The use of a wild type retinas in these co-cultures excluded the possibility of transgene expression in the cultured retina itself, removing the effect of FGF pathway inhibition in that tissue. Furthermore, any GFP expression would be indicative of lens formation only in the transgenic cornea tissue. The entire assembly was then washed through two dishes of culture medium and the recombined eye tissues were cultured individually in 24-well plate wells (1 ml culture medium per well, changed daily). To express the transgene, plates were heat shocked at 34°C for 40 minutes daily in a forced air incubator. Some cultures were not heat shocked for use as controls, and some corneal tissue was derived from larvae that were generated with the control HA-transgenic construct, which upon heat shock expressed only the HA tag. Lens cell differentiation was scored based on the presence or absence of green GFP fluorescence in the cultured eye after 14 days (**Figure 7**). Since those corneas were isolated from transgenic animals, any observed regenerated lenses should express GFP-fused gamma crystalline proteins (Brahma & McDevitt, 1974; Schaefer et al., 1999; Mizuno et al., 1999). These lens-like structures also exhibited a distinct, raised/rounded morphology. Statistical analysis was performed as described below.

**Figure 6:**
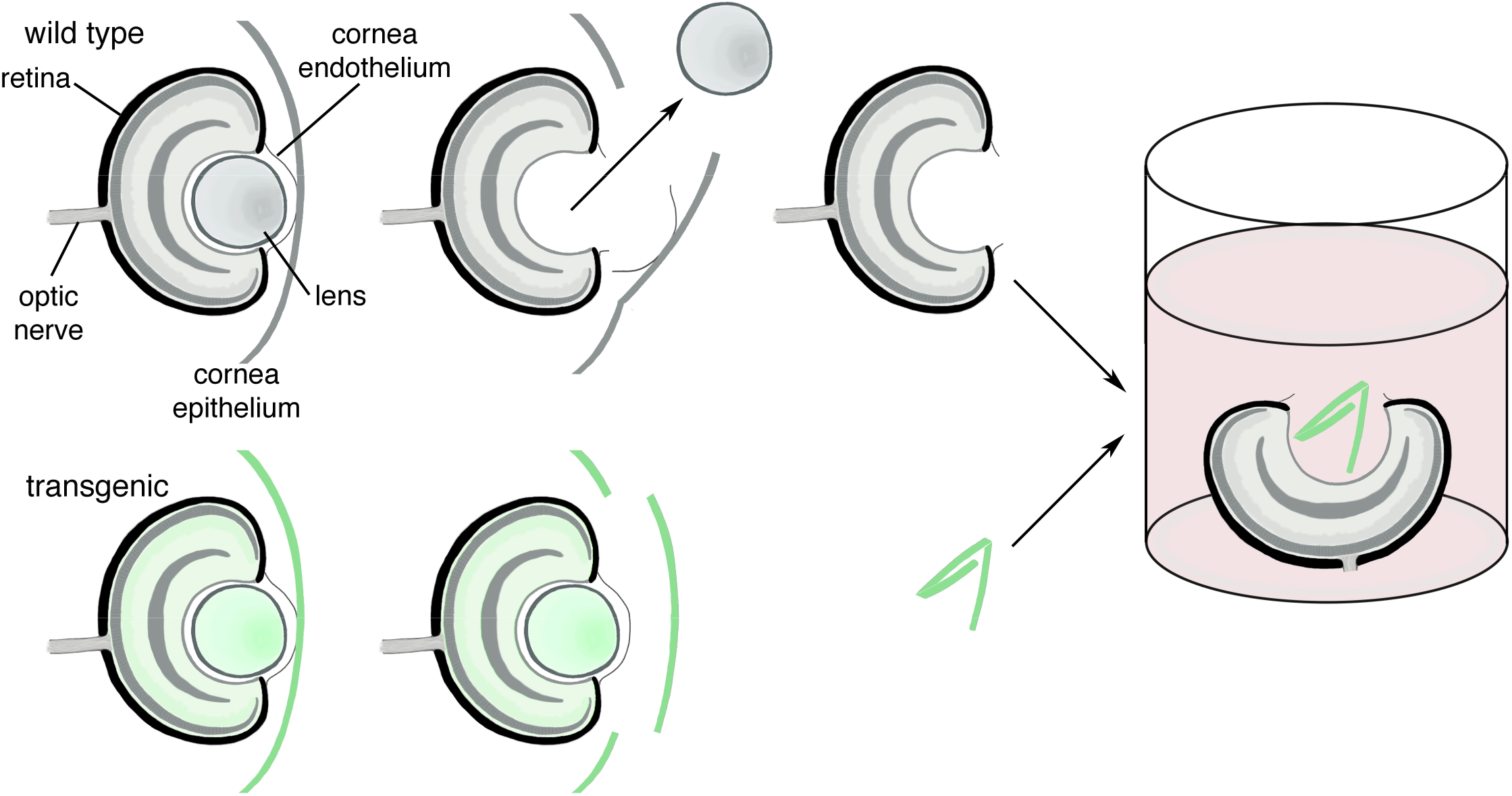
*In vitro* eye culture technique used to assay lens regeneration using transgenic corneas. A wild-type larval eye is shown in the top row, where the lens is removed after incision of the cornea epithelium and endothelium. The cornea epithelium is then removed and the wild-type eye is isolated. The intact transgenic eye is shown on the bottom row, identified here by green color in the lens and eye tissues. The transgenic cornea epithelium is excised from the eye and tucked inside the vitreous chamber of a wild-type eyecup and placed in a culture well. Structures are as labeled.

**Figure 7:**
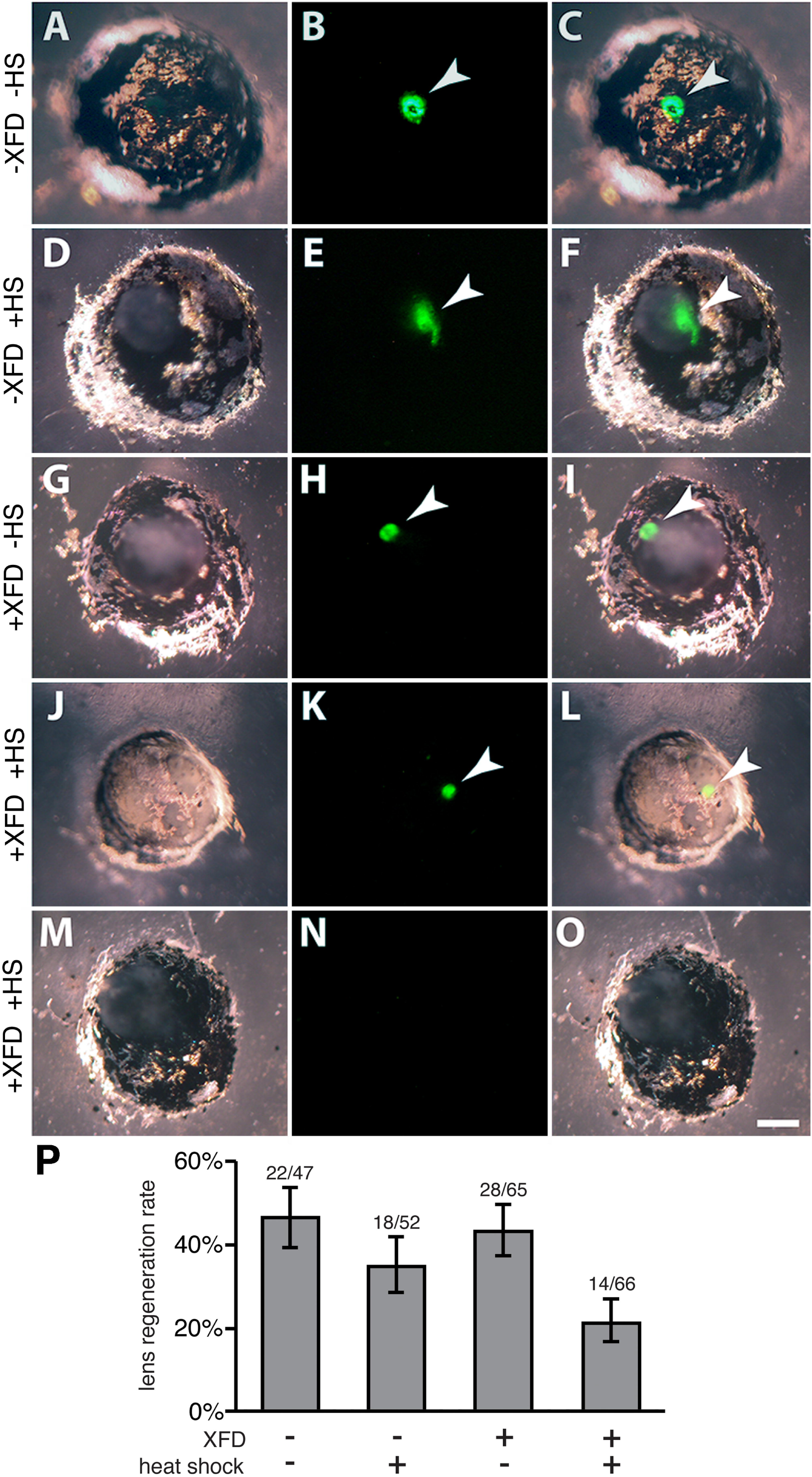
Transgenic corneas cultured in wild-type eye cups for 14 days at room temperature. (A-C) Cultured eye with corneal tissue containing the control HA transgenic construct without XFD and without heat shocks. (D-F) Cultured eye containing the control HA transgenic construct treated with daily heat shocks. (G-I) Cultured eye with XFD transgenic cornea tissue and no heat shocks. (J-O) Cultured eyes containing XFD expressing transgenic cornea, with daily application of heat shocks. (A, D, G, J, M) Cultured eyes under bright field illumination. A lens structure was formed for the case shown in J-L, but not for that shown in M-0. (A, D, G, J, M) Light micrographs of cultured eyes. (B, E, H, K, N) Corresponding fluorescence images of those cultured eyes. (C, F, I, L, O) Corresponding merged images showing both fluorescence and light microscopy images of those eyes. Regenerated lens structures were identified by their green (GFP) fluorescence (arrowheads in B-C, E-F, H-I, K-L). (P) Quantification of lens structure regeneration rates between different transgenic constructs with or without XFD, and either the presence or absence of heat shocks. Lens structure regeneration rates of cultured eyes were assayed after 14 days in culture. The rightmost bar represents the experimental condition in which XFD was expressed in the cornea of the cultured eye. Numbers above each bar indicate the number of cases exhibiting regenerated lens structures as observed for each condition and the total number of eyes cultured for each condition. The Y-axis represents the regeneration rate for each condition. Error bars denote Wilson score confidence intervals with Z=1. The scale bar in O indicates 200 μm.

### Western blotting

XFD expression was verified by Western blot using transgenic larval tails collected after isolation of corneal tissues. Each collected tail was cut in half and each half was placed in a separate 24-well plate well containing culture medium (1mL modified L-15 medium with fetal bovine serum (Kay & Peng, 1991) and antibiotics/antifungals mentioned previously). For each transgenic tail, one cultured half was heat shocked (40 minutes at 34°C in a forced air incubator) and one cultured half was left at room temperature for an equivalent time as a negative control. After heat shock, all tail halves were incubated in culture medium at room temperature for approximately three hours to allow for expression of the transgene. Larval tail halves were flash frozen using a dry ice/ethanol bath approximately three hours after heat shock and stored at -80°C until homogenization for protein expression analysis. Western blot analysis using an anti-HA antibody (sc-805, Santa Cruz) and an anti-rabbit HRP-conjugated secondary antibody was used to confirm XFD expression. A higher molecular weight band (95 kDa) of unknown identity was detected in all samples; however, the band corresponding to XFD was observed only in those transgenic tail halves that were heat shocked (80 kDa, as predicted; **Figure 5G**). Tubulin was also used as a loading control and detected (55kDa) using an anti-beta-Tubulin antibody (T-4026, Sigma-Aldrich, St. Louis, MO).

### Statistics

Statistical analysis to evaluate the significance between experimental and control groups for small sample sizes (n < 40) was performed using Fisher’s exact test, with one-sided p-value calculations. Those samples with a p value less than 0.05 were considered statistically significant. For graphs in **Figures 2** and **3**, Wilson score confidence intervals were used since this method corrects for smaller numbers of trials. Statistical analysis between experimental (XFD) and control groups was performed using the Barnard’s test.

## Results

### FGF signaling is sufficient for lens regeneration in cultured *Xenopus laevis* corneas

FGF1 was shown to be sufficient to induce lens formation in *Xenopus* corneas in a previous study (Bosco et al., 1997b). However, that study had two potential areas for concern. First, only one concentration of FGF1 (500 ng/ml) was examined (Bosco et al., 1997b), which is considered very high when compared to endogenous levels measured in vertebrate eye tissues (typically less than 50 ng/ml in the intraocular space, as determined for human, cow, cat, dog and pig; (Baird et al., 1985a; Caruelle et al., 1989; Tripathi et al., 1992). Second, the culture media used in that study contained 10% fetal calf serum, which likely introduced additional growth factors, including FGFs. A more recent study examined the expression of FGFs in the cornea and retina (Fukui & Henry, 2001). Among the various FGFs examined, FGF1, 8, and 9 mRNAs were differentially expressed in the larval eye, being strongly detected in the retina (Fukui & Henry, 2001). Since the retina releases key factors that support lens regeneration, these three FGFs represented likely candidates involved in triggering lens regeneration. Additionally, we investigated the activity of FGF2 since it has been shown to play an important role in Wolffian lens regeneration in newts and salamanders (Hayashi et al., 2002, 2004, 2006).

Although various cornea culture methods have been designed for mammalian tissue, *Xenopus* cornea culture methods have not been thoroughly developed. The few instances of culturing larval cornea tissues appear to leave the corneas free-floating in culture medium, where they typically form vesicles. In our hands, we found that these cultures tend to lose a significant number of cells (unpublished data, see discussion). This differs from the more flattened epithelial morphology found *in vivo*. Here, we employed a novel *in vitro* cornea culture method to assay various factors for *Xenopus* lens regeneration that allows the corneas to attach and spread on the bottom surface of gelatin-coated plastic tissue culture wells, and these cultures appear to spread and do not lose cell mass (**Figure 1**).

To ensure that the FGF proteins were active and effective in our cultured cornea experiments, a protein activity assay was performed on cultured *Xenopus* eyes. Following the addition of FGF protein, the eyes were fixed and corneas were removed and assayed for increased cell activity in the cornea, which was observed by labeling chromatin with anti-Histone H3S10P antibody (**Figure 4**). The observed results suggest that the addition of FGF1, 2, and 8 in eye cultures resulted in increased chromatin labeling in the cornea when compared to control corneas. Activity was confirmed for FGF1 (25% increase in histone labeled cells), FGF2 (39% increase in histone labeled cells), and FGF8 (11% increase in histone labeled cells). FGF9 failed to induce an enhanced proliferative response in these cornea tissues (no increase observed above control corneas). The negative results obtained for FGF9 were verified with a second, independent sample of FGF9 in a second trial of experiments, with the same result (data not shown).

Next, we observed the effect of a range of concentrations of FGFs on *in vitro* cultured corneas in minimal medium (67% L-15 medium without added serum, as serum may contain various growth factors, including FGFs). Control cultures without FGF protein were also evaluated. As shown in **Figures 1** and **2**, cultured corneas quickly attach and spread on the bottom of their respective wells. After 9 days in culture, corneas were evaluated for regenerated lenses using anti-lens antibody staining (Henry & Grainger, 1990). This antibody has been used in numerous previous studies (Henry and Grainger, 1990; Fukui & Henry, 2001; Walter et al., 2008; Perry et al., 2010, 2013) which is ample time to observe any lens cell formation (Henry & Mittleman, 1995). In addition to positive anti-lens staining, the small lenses had a raised, three-dimensional, rounded morphology as opposed to the flattened epithelial appearance of the surrounding corneal tissue.

Small lenses were observed following FGF1 treatment at all concentrations (**Figure 2**). A statistically significant increase in lens formation was observed with an FGF1 concentration as low as 50 ng/ml relative to FGF (-) controls (p = 0.011; **Figure 2**). Higher concentrations (100 ng/ml FGF1 and above), resulted in a more gradual increase in the rates of lens formation. However, when comparing regeneration rates at 100 ng/ml of FGF1 versus 1 ug/ml of FGF1, the slight increase observed over this range of concentrations was not statistically significant (p = 0.36 comparing the 100 ng/ml and 1 μg/ml conditions). The EC_50_ for FGF1 application appears to be between 20 ng/ml (19% regenerated) and 50 ng/ml (26% regenerated), assuming a maximal lens regeneration rate of approximately 50% at the highest FGF1 concentrations, which is similar to the rates observed *in vivo*. From these observations of regeneration rates, we can conclude that FGF1 application is sufficient to trigger lens formation, and effective at concentrations as low as 50 ng/ml FGF1.

FGF8 was also tested over a similar range of concentrations (10, 20, 50, 100, or 500 ng/ml FGF8). There were few cases of positive anti-lens staining observed (**Figure 3**). No cases of regeneration were observed for concentrations of FGF8 below 100ng/ml (**Figure 3**). Higher concentrations of FGF8 resulted in some lens regeneration in 1/15 cases (100 ng/ml) and in 3/16 cases (500 ng/ml, p = 0.2258). These results indicate that FGF8 has a very limited potential to induce lens regeneration and only at concentrations above 100ng/ml.

**Figure 3:**
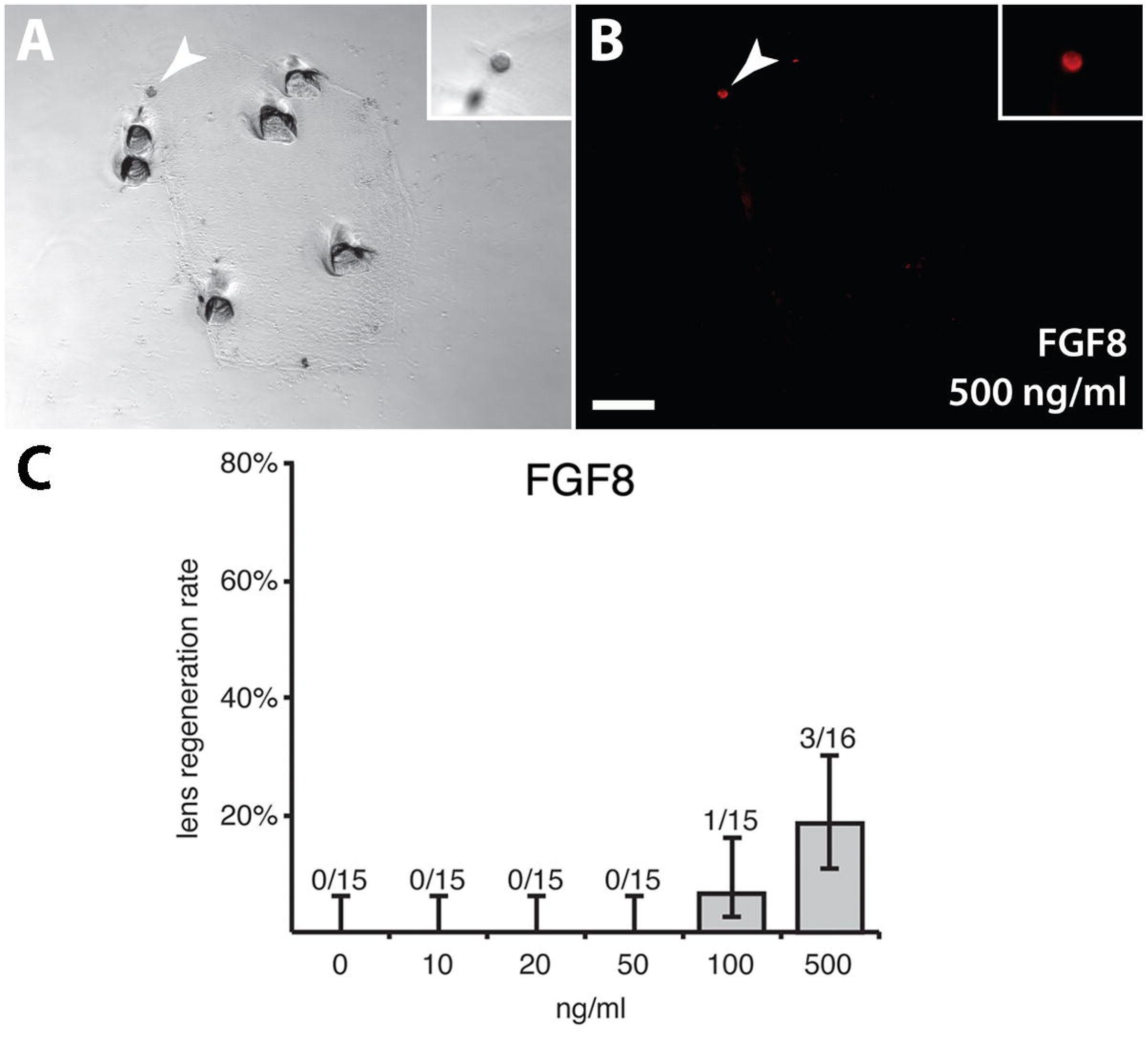
Anti-lens antibody labeling of FGF8 treated cultured cornea (9 days). This cornea was treated with 500 ng/ml FGF8 (A-B). Images were taken under bright field (A) and fluorescence illumination to detect anti-lens antibody staining (B). The inset in A-B contains a higher magnitude image of the correspond lens in each panel. White arrowheads in A-B point to regenerated lens structures. Depressions in the culture well (black circular spots in A) indicate points (indentations) where the cornea was secured to the bottom of the culture well using sharpened forceps. The scale bar in B indicates 100 μm for the insets in A-B and 200 μm for panels A-B. (C) Quantification of lens structure formation rates for FGF8. The leftmost bar (0 ng/ml) indicates the control condition, where no FGF was applied. The number of corneas forming lens cell aggregates and total number of corneas observed for each condition are indicated above each bar. The Y-axis indicates the % regeneration rate for each condition. Error bars show Wilson score confidence intervals where Z=1.

A range of concentrations for FGF2 and FGF9 (up to 500 ng/ml) were also tested for their potential to induce lens regeneration (**Table 1**). Very few cases of lens regeneration were observed, with no condition exhibiting significantly more lens regeneration when compared to FGF (-) controls. Thus, FGF2 and FGF9 do not appear to be sufficient for inducing lens regeneration in these cultured corneas. The results of the protein activity assay for FGF9 also showed no increase in histone H3 in the cornea following addition of the protein, further suggesting that FGF9 does not induce a response in these tissues. Previous studies indicate that this protein plays roles mainly in the nervous system and sex determination (Miyamoto et al., 1993; Kim et al., 2006). On the other hand, FGF2 was shown to stimulate proliferation, yet it had no activity as far as stimulating lens regeneration in the cornea. Together, these experiments suggest that FGF1 is a potent inducer of lens regeneration and FGF8 may have a limited capacity to induce lens formation.

### XFD expression inhibits lens regeneration

Lens regeneration was inhibited upon expression of the dominant negative FGF receptor (XFD) in corneal tissue, assayed using an *in vitro* culture method with transgenic larval eye tissue as described in the Experimental Procedures section (**Figures 5-7**; Fukui & Henry, 2001; Thomas & Henry, 2014). Only 14/66 eyes (21%) regenerated lenses under the experimental condition, in which the XFD transgene was expressed under daily heat shock (**Figures 5, 7**). This rate of regeneration was lower than that seen in the three control conditions examined. In the first control set, wild-type eye cups were cultured with transgenic corneas with the control HA transgene (with the potential to express only the HA tag; see **Figures 5, 7**) and with no application of heat shock. These cultures regenerated lenses in 22/47 cases (47%; p = 0.0042; **Figure 7**). As a second control, eye cups were combined with transgenic corneas with the control HA transgene, and heat shocked on a daily basis. These cultures regenerated lenses in 18/52 eyes (35%; p = 0.1180; **Figure 7**). In the final control, transgenic XFD corneas implanted into wild-type eye cups but were not exposed to heat shock (thus not expected to express XFD). These cases regenerated lenses in 28/65 eyes (43%; p = 0.0074; **Figure 7**). Among the three control conditions, lens regeneration does not differ significantly: all pairwise comparisons between these regeneration rates have p > 0.1. Although heat shock itself may somewhat inhibit lens regeneration, it does not significantly affect the rate of regeneration compared to wild-type controls without heat shock. However, XFD expression in conjunction with heat shock appears to inhibit this process, indicating a role for FGFR signaling in lens regeneration.

## Discussion

### FGF1 strongly induces lens formation in cultured corneas

Using a cornea culture assay, we have shown that FGF1 is sufficient to trigger lens formation. This is in agreement with one study (Bosco et al., 1997b) showing FGF1 can trigger lens regeneration. Here, however, we examined a range of FGF1 concentrations and used a different approach in culturing *Xenopus* corneas. Lenses were observed as early as 5 days in culture, consistent with that previous study (Bosco et al., 1997b). Our study finds that FGF1 concentrations as low as 50 ng/ml are sufficient for inducing lens regeneration. These concentrations are an order of magnitude smaller than the single concentration examined by Bosco et al. (1997b; 500ng/ml). We also observed that FGF8 could be a weak inducer of *Xenopus* lens regeneration, though the differences were not found to be all that significant. FGF9 and FGF2 do not appear to significantly enhance lens regeneration at all. Overall, these experiments show that FGF1 is a potent inducer of lens regeneration.

One possible explanation for FGF8 triggering some lens formation at high concentrations may be that FGF8 shares affinity for several FGFR isoforms with FGF1 (Ornitz and Itoh 2015), potentially triggering the same FGFR signaling cascade that prompts lens regeneration. FGF2 can also activate the same FGFR isoforms that FGF8 activates, potentially activating the same FGFR downstream signaling pathways (Ornitz and Itoh 2015); however, FGF2 did not trigger lens formation in our assay.

A notable difference between Wolffian lens regeneration and our system concerns the specific FGF ligand sufficient for lens formation. Our results indicate that FGF1 specifically induces the formation of lens cell aggregates in cultured *Xenopus* corneas. In the newt, FGF1 is first detectable and becomes upregulated in the regenerating lens vesicle, and subsequent lens epithelium and lens fibers beginning at day 10 of the regeneration process (Del Rio-Tsonis, 1997). Yang et al. (2005) investigated the ability of the newt to regenerate a lens following both neural retina and lens removal either with or without the addition of rnFGF-1. Under these conditions parts of the retinal reform and both the dorsal and ventral iris underwent depigmentation/dedifferentiation to produce a regenerated lens, but those regenerated lenses were characterized by abnormal fiber cells and a thin or absent lens epithelium. Their results suggest that the timing of FGF1 expression could be important for proper Wolffian regeneration.

In contrast, FGF2 appears to be a key factor to trigger Wolffian lens regeneration (Hayashi et al., 2002, 2004), but it is not able to induce lens regeneration in *Xenopus*. Therefore, FGF1 and FGF2 likely do not share a conserved deployment during the two processes of lens regeneration. These findings add to a growing base of knowledge showing various forms of lens regeneration employ different signaling mechanisms and evolved independently (e.g. retinoic acid and Wnt signaling; Hamilton et al., 2016; Thomas & Henry 2014).

Though we have not measured the concentration of FGF proteins in *Xenopus* eyes, we show that the lower FGF1 concentrations sufficient to trigger lens cell differentiation in cultured *Xenopus* cornea tissue reflects more similarly with endogenous FGF levels, as measured in other vertebrate eyes (e.g., human, cow, cat, dog and pig; Baird et al., 1985a,b; Tripathi et al., 1992). For example, FGF1 is expressed more highly in the bovine retina than the cornea: 35 ng/ml FGF1 has been measured in the aqueous humor (Caruelle et al., 1989). We detected less *FGF1* mRNA in the *Xenopus* cornea than the retina (Fukui & Henry 2001). Furthermore, approximately 1ng/ml of FGF2 is present in the aqueous humor of humans, cats, dogs, and pigs (Tripathi et al., 1992), and human intraocular FGF2 levels may be around 10-20 ng/ml (Baird et al., 1985a). In rat lens epithelial cell cultures, the levels of FGF2 required for inducing lens fiber differentiation are 40 ng/ml (Lovicu & McAvoy 1989; McAvoy & Chamberlain 1989).

A potential limitation of our study lies in the use of human recombinant FGFs to treat *Xenopus* tissues. Although these are highly conserved proteins, there are some differences in composition between human and *Xenopus* FGFs and FGFRs, including the FGF-FGFR binding sites (Plotnikov et al., 2000; Stauber et al., 2000). It may be that the FGFs obtained from these different sources have different potencies.

The response of *Xenopus* corneal tissues to FGF1 described here may explain the results of previous xenoplastic grafting experiments in which eyecups from other frog species (incapable of lens regeneration) were implanted with *Xenopus* corneas that subsequently exhibited lens regeneration (Filoni et al., 1979; Bosco et al., 1991, 1993; Henry & Elkins 2001). It is likely that the formation of lenses in these studies indicates that all of those eyecups supply FGF1.

### The potential effects of morphology in cornea cultures

Our culture scheme for *Xenopus* permitted corneas to attach and spread on the bottom of the culture well. Although a previous study (Bosco et al., 1997b) utilized cultured *Xenopus* corneas, those free-floating corneas did not attach to any surface and became vesicularized (**Figure 1**). This was similar to an early investigation of cultured *Xenopus* corneas that reported hollow vesicle formation in cases where the cornea tissue was not attached to a substrate (Campbell & Jones, 1968). In contrast, our method is more similar to many mammalian cornea epithelial culture models, which are typically attached to a coated surface to form an epithelial sheet (Reichl et al., 2011; Sabater et al., 2013), with the exception of a few studies on sphere-forming cultures (Mimura et al., 2005; Uchida et al., 2005; Yokoo et al., 2005). Normally the inner surface of the cornea epithelium has a specialized basement membrane (Bowman’s layer), which may be mimicked in our cultures by attachment to the gelatin-coated substrate. Notably, we observed that the lens structures that appeared tended to be localized to the periphery of the corneal cultures. Regenerated lenses arise from the basal epithelial cells *in vivo* and this layer lies closest to the substrate in these culture experiments, which could limit exposure to soluble FGFs. The presence of lens structures at the periphery could be due to greater exposure to FGFs. Nonetheless, one can argue that the method presented here tends to maintain a more natural epithelial morphology *in vivo*.

This culture method results in distinct lens-like structures forming within the cultured corneas following the addition of FGF1 (**Figure 2**). These lens structures have a three-dimensional rounded appearance distinct from the flattened epithelial appearance of the remaining cultured cornea tissue. In contrast, Bosco et al. (1997b) found that following FGF treatment, the entire free floating cultured cornea appears to stain positively for lens proteins after 7-10 days. The differences in culture methods may contribute to differences in cornea sensitivity to lens regeneration factors *in vitro.* The flattened epithelial morphology could promote diffusion and favor the regeneration of isolated lens structures. In both cases, the resulting lens structures appear to be similar in size (on the order of 100μm in diameter). This implies that either there was extensive cell loss/death in the vesicular cultures of Bosco et al. (1997b), or the amount of cornea tissue that was initially isolated in those experiments (not described) was much smaller than the intact corneas that were isolated here. In fact, preliminary attempts to replicate the experiments by Bosco et al. (1997b) resulted in extensive cell death/loss where only a small central vesicular structure remained, which raised concerns about the integrity of those free floating cultures (data not shown). Given past insights into tissue and cell morphology affecting cell proliferation (Watt et al., 1988), survival (Folkman & Greenspan, 1975; Chen et al., 1997), signaling (Rangamani et al., 2013), and differentiation (Kumar et al., 2011; Nampe & Tsutsui, 2013), it is likely that morphology may also play a key role in *Xenopus* lens regeneration.

### FGFR signaling is necessary for lens regeneration

It has been established that the regenerated lens originates from the inner layer of the cornea epithelium in larval *Xenopus laevis* (Freeman, 1963). Previous studies indicated that key signals from the neural retina are required to trigger lens regeneration in *Xenopus* (Filoni et al., 1981, 1982, 1983, 1993c; Reeve & Wild, 1981, Bosco & Filoni, 1992; Bosco et al., 1997b). Here we tested the necessity of FGFR activity by expressing a dominant negative FGFR1 transgene known to inhibit the activation of various endogenous FGFRs (Amaya et al., 1991). The *Xenopus laevis* dominant negative FGFR1 transgene (XFD) was initially developed to study *Xenopus* mesoderm development during early embryogenesis (Amaya et al., 1991, 1993). XFD acts by heterodimerizing with endogenous FGFRs to inhibit downstream activation of the pathway (Amaya et al., 1991). By examining transgenic corneas cultured within non-transgenic eye cups, we found that FGFR function, specifically in corneal tissue, is important for lens regeneration. This demonstrates a more specific necessity of FGF signaling in lens regeneration than the experiments using SU5402 to inhibit FGFR function (which may also inhibit VEGFR function; Sun et al., 1999; Fukui & Henry, 2001).

At this point, we cannot distinguish specifically whether FGFR1 is necessary, as opposed to other FGFRs, because the dominant negative FGFR1 can heterodimerize and inhibit the functions of FGFR1, FGFR2, and FGFR3 (Ueno et al., 1992). In fact, all three of these receptors (but not FGFR4) are expressed in *Xenopus* cornea epithelial tissue (Fukui & Henry 2001). There is some evidence that expression of different dominant negative FGFRs may have slightly different developmental effects (Hongo et al., 1999; Ota et al., 2009). Therefore, future experiments could investigate the effects of other dominant negative FGFRs, on lens regeneration in this system. These findings are consistent with a previous correlation observed between FGFR2 expression and competence to form a lens in *Xenopus* (Arresta et al., 2005). Tissues that are competent to regenerate a lens include only the cornea and pericorneal ectoderm, (Freeman, 1963; Bosco et al., 1979; Cannata et al., 2003; Arresta et al., 2005) and Arresta et al., (2005) showed that only those lens competent epithelial tissues expressed FGFR2 (the “Bek” isoform).

Although inhibition of FGFR function caused reduced rates of lens regeneration, not all regeneration was inhibited upon XFD expression. There are some potential explanations for this. F0 animals generated using the REMI transgenesis method are known to have different copy numbers of the transgene inserted into the genome (Kroll & Amaya, 1996; Sparrow et al., 2000). Therefore, there may be a difference in copy numbers of the transgene for each larval cornea tested, affecting the strength of FGFR inhibition. In addition, some mosaicism in expression within individual larvae has been observed using this transgenesis method (Sparrow et al., 2000), which raises the possibility that some transgenic larvae did not express the transgene in every cornea cell. Nevertheless, our experiments show a substantial decrease in lens regeneration rates upon XFD expression.

## Acknowledgments

We thank Dr. Peter Jones (University of Massachusetts) for the CGHG plasmid and technical assistance regarding the transgenesis protocol. We thank Dr. Richard Harland (University of California, Berkeley) and Dr. Enrique Amaya (University of Manchester, UK) for supplying the plasmid containing XFD (Sive et al., 2000). Finally, we thank Alvin Thomas and Paul Hamilton for help in editing this manuscript and figures. This work was supported by NIH grants EY09844 and EY023979 to JJH.

